# Prioritising Functional Noncoding Variants via eRNA Post-transcriptional Interaction Maps in Human Samples

**DOI:** 10.1101/2025.07.31.667983

**Authors:** Natalia Benova, Rene Kuklinkova, Jessica L. Haigh, James R. Boyne, Chinedu A. Anene

## Abstract

Noncoding variants and mutations outnumber their coding counterparts but remain challenging to interpret functionally. We present TranCi, a method that prioritises human genetic variations by integrating enhancer RNA (eRNA) expression with eRNA-mRNA interactome maps. By linking variant-associated changes in eRNA to downstream gene regulation, TranCi captures functional effects missed by sequence-based or chromatin-centric approaches. In esophageal squamous cell cancer, TranCi identifies noncoding mutations with roles in disease initiation and progression. A personalised mode enables analysis at single-patient resolution, uncovering potential individual-specific regulatory variants. TranCi thus provides a mechanistic framework for interpreting noncoding variations and uniquely identifies their downstream targets, where standard methods often fall short. TranCi is available as a module within the eRNAkit R package (https://github.com/AneneLab/eRNAkit), leveraging its database of eRNA expression and interactions for functional variant interpretation.

## Introduction

Noncoding variants in the human genome outnumber those in coding regions by orders of magnitude, yet their functional interpretation remains a major challenge in human genomics ^1,2^. While large-scale initiatives such as TCGA and ICGC for cancer, along with TOPMed and NHLBI for cardiovascular traits, PsychENCODE for neuropsychiatric disorders, and AMP-AD for Alzheimer’s disease, have unveiled a rich catalogue of genetic variants, only a minority have been conclusively linked to regulatory functions ^3–7^. Among these, most validated impacts are still linked to target genes through coding and their proximal regions (i.e., promoters and UTR), despite these regions representing only a small fraction of the total mutational burden in diseases such as cancer. Complementary efforts such as ENCODE, Roadmap Epigenomics, and GTEx have expanded our understanding of the noncoding genome, revealing a vast landscape of regulatory elements whose perturbation may contribute to disease susceptibility and phenotypic diversity ^8–10^. However, elucidating the regulatory consequences of variations on these elements has proven difficult due to the lack of reliable strategies for linking distal mutations to their target gene functions. Enhancer RNAs (eRNAs), transcribed from active enhancers, offer a mechanistically informative layer for interpreting noncoding variation. Unlike static sequence features, eRNAs reflect real-time regulatory activity and can engage in cis- and trans-interactions that influence mRNA stability, translation, and transcriptional output.

Traditional approaches to noncoding variant interpretation rely on chromatin features (e.g., TADs, histone marks) or sequence motifs, but these offer limited resolution: chromatin contacts span broad regions, and motif hits in distal regulatory elements lack functional validation. Variant scoring tools like RegulomeDB and DeepSEA ^11,12^ assess potential regulatory activity based on chromatin or sequence features but do not account for the downstream targets and biological impact of the variations. eRNAs offer a unique opportunity to bridge this gap. Initially characterised as markers of enhancer activation, eRNAs are now recognised as active molecules that engage in diverse cis-and trans-regulatory functions, including facilitating enhancer– promoter communication, promoting chromatin remodelling, recruiting transcriptional co-activators, sequestering repressive factors, and engaging in RNA–RNA and RNA–protein interactions that modulate target mRNA stability and translation ^13–19^.

These context-dependent roles position eRNAs as both sensors and effectors of regulatory activity, capable of transmitting the impact of noncoding mutations to target gene expression. By integrating eRNA dynamics with somatic variant data, it becomes possible to mechanistically link distal enhancer mutations to their functional outcomes.

To this end, we developed TranCi, a computational framework that integrates noncoding variations with eRNA expression and RNA–RNA interactome maps to prioritise those with potential regulatory impact across disease contexts. TranCi identifies variants overlapping eRNA loci, tests for differential eRNA expression between variant carriers and non-carriers and connects dysregulated eRNAs to their target mRNAs via experimentally derived RNA–RNA interaction networks (Figure 1). A composite scoring scheme integrates evidence from eRNA expression, RNA–RNA interactions, and downstream mRNA impact to prioritise variants as empirical, predictive, or weak, reflecting increasing functional support. This provides mechanistic insight into the regulatory impact of noncoding variations, without relying on chromatin proximity-derived inference alone.

**Figure 1.**
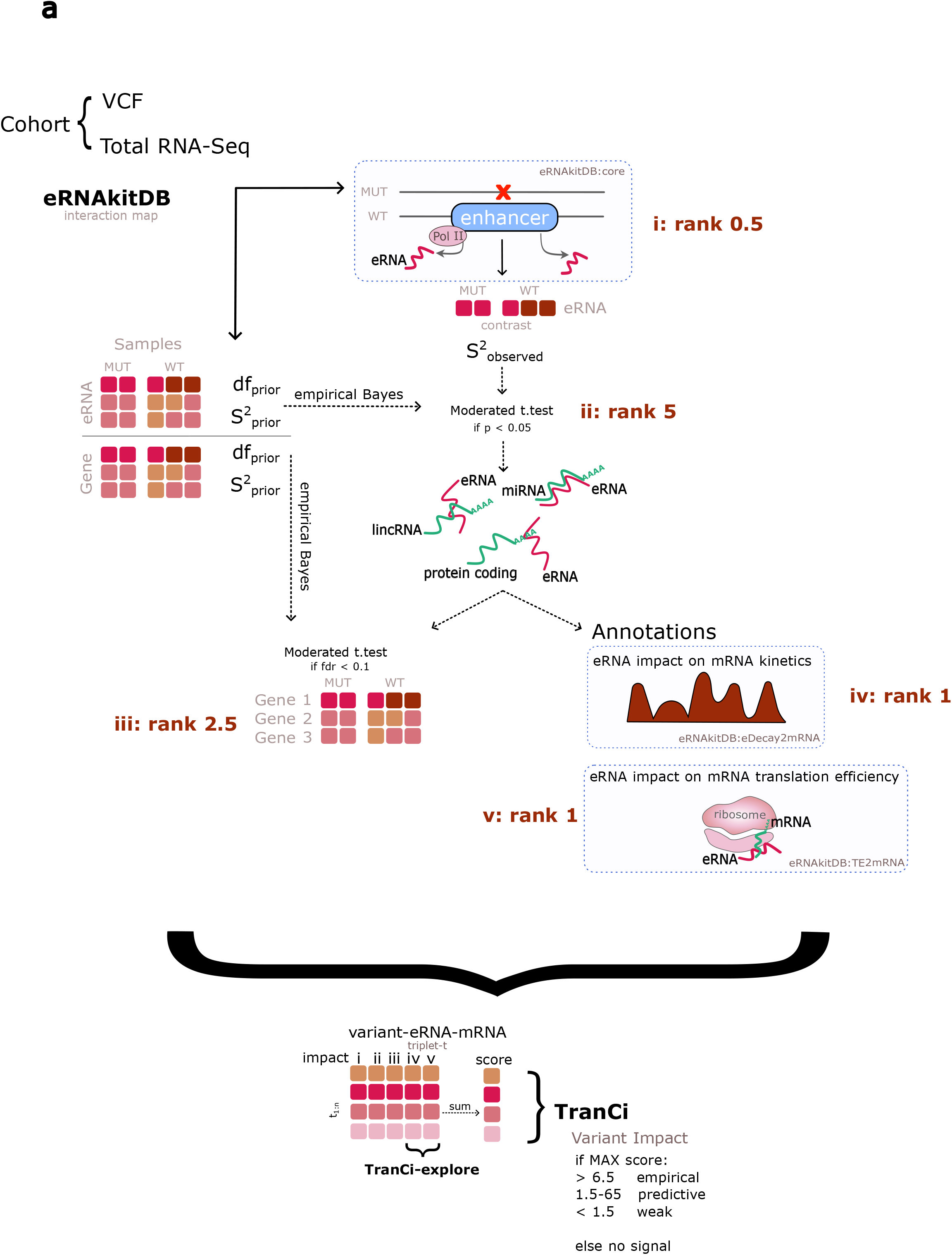
TranCi Framework. a) TranCi is a computational framework to prioritise regulatory noncoding variants by integrating eRNA expression and RNA-RNA interaction data. First, samples are grouped based on variant presence or absence. Variants are mapped to enhancer RNA (eRNA) loci, and differential expression analysis is performed to assess variant effects on eRNA activity. Next, eRNAs are linked to their mRNA targets through RNA-RNA interaction networks, enabling testing of variant impact on target gene expression. Subsequently, TranCi incorporates previously derived eRNA effects on mRNA stability and translation efficiency to further characterise variant consequences. A composite scoring scheme then integrates these evidence layers to prioritise variants with functional regulatory potential. b) TranCi-explore extends he core functionality to personal genome context.

An important aspect of TranCi is its ability to detect functional signals even with sparse genotype stratification. Since noncoding variants are often rare, traditional differential expression tests can suffer from low statistical power. To overcome this, TranCi employs an empirical Bayes moderated t-statistics approach to stabilise variance estimates by borrowing strength across all measured transcripts. This variance shrinkage strategy enhances sensitivity while controlling false discovery rates, enabling robust detection of variant-associated expression changes in small patient subgroups. We demonstrate TranCi by applying it to whole-genome and transcriptome data from esophageal squamous cell carcinoma (ESCC), identifying novel noncoding variants that disrupt eRNA function and impact oncogenic networks, many of which are missed by existing tools. TranCi-explore further extends this capability to individual genomes, enabling personalised variant interpretation and advancing translational genomics.

## Results

### Core Database Supporting TranCi Functional Analysis

To support mechanistic interpretation of noncoding variant via eRNA activity, we built TranCi on top of eRNAkitDB, a curated database integrating eRNA annotations, expression profiles, RNA–RNA interactions, 3D chromatin contacts, and regulatory associations across diverse human cell types ^18^. This multi-layered resource provides the molecular context needed to link genetic variations to dysregulated eRNA activity and downstream gene targets, enabling TranCi to systematically prioritise functional distal regulatory variants in disease. At the core is the RNA–RNA interaction (R2R) dataset, which, together with annotation of 3D genome contacts, supports variant mapping to eRNA–mRNA pairs. Functional interpretation is further refined by incorporating data on eRNA-driven effects on mRNA stability and translation ^19^. Together, these components allow TranCi to connect sequence-level perturbations to gene regulatory outcomes with greater specificity and resolution (Figure 1).

### Noncoding eRNA-associated variants reveal hidden regulatory landscapes in ESCC

To evaluate the utility of TranCi in a disease context, we applied the framework to esophageal squamous cell carcinoma (ESCC), a genomically well-characterised cancer with a high frequency of somatic mutations, particularly in TP53 and a small set of other protein-coding genes ^20^. While several studies have begun to explore the functional significance of noncoding mutations in ESCC development and progression, these efforts have largely been restricted to promoter regions, untranslated regions (UTRs), and annotated long noncoding RNAs (lncRNAs) ^1,21,22^. As such, large portions of the noncoding genome, particularly distal enhancer regions, remain unexamined. The availability of whole-genome sequencing and matched total RNA expression data in TCGA makes ESCC a compelling test case for TranCi, which extends the search for functional somatic variants into enhancer RNA (eRNA) loci and their potential regulatory targets.

We applied TranCi framework to 66 TCGA-ESCC samples with matched total RNA-Seq and whole-genome somatic noncoding variant calls using the “TranCi()” function in eRNAkit. Variants were first filtered by recurrence (≥2 samples) as specified by the -*n* parameter, and those overlapping curated eRNA loci were retained for further analysis. In total, TranCi identified 27 empirical, 380 predictive, and 675 weak regulatory variants in this cohort (Table S1-S2). The empirical variants were distributed across multiple chromosomes, including, 1:215324880:A:T, 1:39925188:C:G, 1:39925563:C:A, 1:39925714:C:G, 1:39925935:C:T,

1:39925967:C:T, 3:125218451:G:C, 3:131425605:T:C, 3:133914588:A:G,

3:180350156:G:A, 3:75279980:A:G, 4:59335452:C:A, 4:62691042:C:G,

5:171805734:G:A, 5:171835823:G:T, 6:150306716:T:G, 8:48761223:A:G,

8:56885579:A:G, 8:81945852:A:C, 8:92370027:G:T, 8:96177928:A:G,

12:94715822:A:G, 18:72947422:G:T, 18:72977901:C:A, 19:28140279:T:C,

19:28187045:G:T, 19:28190674:T:C. Among these, only one variant, 1:215324880:A:T, was previously annotated in public databases (dbSNP : rs185317019).

### TranCi identifies regulatory noncoding variants missed by chromatin-based models

To evaluate how TranCi’s variant prioritisation compares with existing tools, we benchmarked its classifications in ESCC against those from RegulomeDB and DeepSEA ^11,12^, two widely used methods for assessing regulatory potential of noncoding variants. Given that TranCi integrates eRNA expression and RNA–RNA interaction data, we hypothesised it would identify functional noncoding variants missed by sequence- or chromatin-based approaches.

To evaluate the concordance between TranCi variant classification and RegulomeDB rankings, we performed a chi-squared test comparing the distribution of RegulomeDB categories across TranCi-defined variant classes (empirical, predictive, and weak). The result was non-significant (χ^2^ = 13.783, df = 14, p = 0.466, Figure 2b), indicating no strong statistical association between the two methods. Despite this lack of concordance, a closer inspection reveals important insights. Among the 27 empirical variants, those supported by both genotype–expression associations and eRNA–mRNA interactions, none were ranked in the top RegulomeDB categories (i.e., scores 1f–3a). Instead, most fell into lower-confidence ranks (scores 4–7), which typically lack chromatin or motif support. Similarly, the 378 predictive variants, which had enhancer–target support but no genotype correlation, were largely assigned modest or low RegulomeDB scores. These patterns suggest that RegulomeDB fails to prioritise variants that TranCi identifies through eRNA-informed evidence. Interestingly, variants in the weak class, those lacking either empirical support, were more often annotated with higher RegulomeDB scores (Ranks 2a and 3a, n=29, Figure 2a), reflecting chromatin features such as transcription factor binding or DNase sensitivity. This alignment with chromatin signatures, but not with transcriptional evidence, reinforces that TranCi prioritises variants based on regulatory function rather than static sequence context.

**Figure 2.**
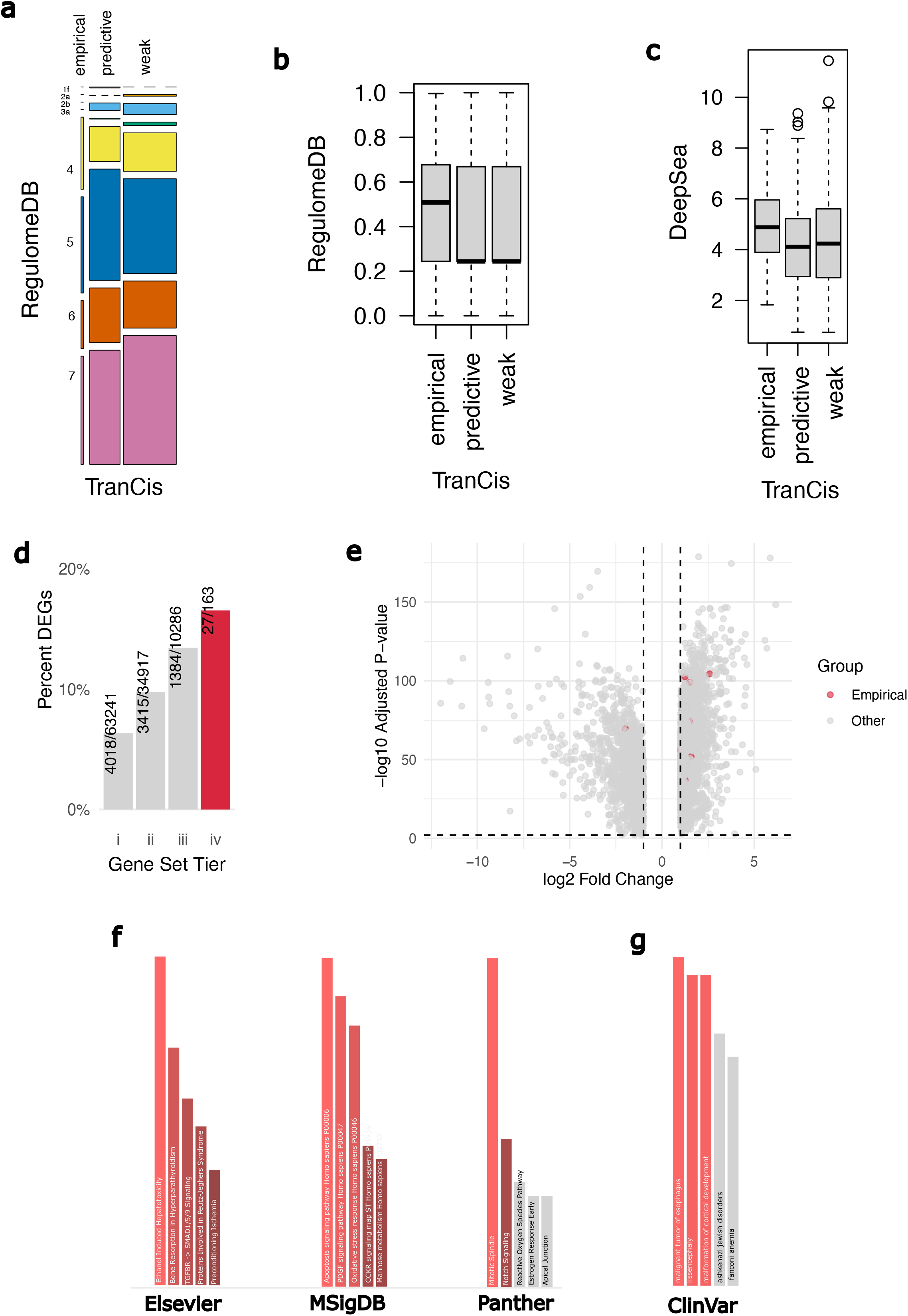
TranCi captures orthogonal regulatory signals not reflect existing annotation frameworks. (a) Cross-classification of TranCi-prioritised noncoding variants and RegulomeDB classes. (b) RegulomeDB probabilities stratified by TranCi classes. (c) Summed DeepSEA probabilities stratified by TranCi classes. (d) Proportion of differentially expressed genes (DEGs) across TranCi prioritisation tiers: (i) all annotated genes in the transcriptome, (ii) genes with eRNA-mRNA interactions in the R2R annotation, (iii) genes associated with TranCi-prioritised variants (weak to empirical), and (iv) genes in the empirical triplets. (e) Volcano plot of all DEGs in ESCC compared to normal Esophageal tissue (GEPIA2 – esca), with TranCi-associated empirical targets highlighted in red. (f-g) Pathway enrichment (Enrichr) of TranCi empirical target genes in (f) Elsevier Pathway Collection, MSigDB Hallmark 2020, Panther 2016, and (g) ClinVar 2019.

In addition to categorical ranks, RegulomeDB provides a probability score that reflects the likelihood of a variant having regulatory function based on integrated epigenomic and motif data. While these scores are correlated with rank, they offer a continuous metric for finer comparisons. To assess whether TranCi variant classes differ in their RegulomeDB probability distributions, we applied a Kruskal–Walli’s rank sum test. The result was not statistically significant (χ^2^ = 3.1433, df = 2, p = 0.2077, Figure 2b), indicating no meaningful difference in RegulomeDB-derived regulatory probabilities across TranCi’s empirical, predictive, and weak classes. This lack of differentiation further supports the notion that TranCi prioritises a distinct set of functional noncoding variants, relying on transcriptional and interaction evidence that may not be captured by sequence-or chromatin-derived annotations alone.

To further evaluate this distinction, we benchmarked TranCi variant classes against DeepSEA, a deep learning model trained to predict the chromatin effects of noncoding variants across hundreds of epigenomic marks. For each variant, a single composite DeepSEA score was computed as the sum of all predicted probabilities from the model, representing cumulative chromatin impact across all features. While empirical TranCi variants tended to show slightly higher DeepSEA scores overall, the difference across classes was not statistically significant (Kruskal–Wallis χ^2^ = 5.026, df = 2, p = 0.08103, Figure 2c). Many highly ranked variants had only moderate or low DeepSEA predictions, further suggesting that TranCi identifies a regulatory axis orthogonal to conventional sequence-and chromatin-based models.

### TranCi’s high-confidence regulatory variants are enriched for chromatin contact, but not dependent on it

Across all the variant–eRNA–mRNA triplets evaluated in the final scoring, only 1.408% (550 out of 39,058) showed direct evidence of chromatin interaction within topologically associating domains (TADs) between the eRNA and it’s target mRNAs ^18^. When grouped by integrative scores into “weak” (<1.5), “predictive” (1.5–6.5), and “empirical” (≥6.5) tiers, a clear trend emerged: 10.84% of empirical triplets had TAD-level support, compared to just 0.3–1.62% in the lower tiers. This enrichment supports the use of high TranCi scores as markers of biologically plausible enhancer–target relationships. Importantly, most high-scoring predictions, including more than half of the empirical tier, lacked known 3D genome interaction evidence. This highlights the key strength of the TranCi framework: by leveraging RNA-RNA interaction signals between eRNAs and their targets, it uncovers putative regulatory links that may operate independently of chromatin proximity, offering a complementary and potentially novel layer of functional annotation.

### Noncoding variants identified by TranCi converge on oncogenic pathways and cancer dependencies in ESCC

To explore whether the predicted target genes of high-scoring (empirical) variant– eRNA–mRNA triplets converge on specific biological pathways, particularly those relevant to ESCC, we performed a multi-layered functional characterisation.

### High-priority regulatory variants are linked to differentially expressed genes in ESCC

We first assessed whether the putative mRNA targets of these empirical variants were more likely to be differentially expressed in esophageal carcinoma compared to normal esophageal tissues. We established a four-tiered gene set system: (i) all annotated genes in the transcriptome, (ii) genes with eRNA-mRNA interactions in the R2R annotation, (iii) genes associated with TranCi-prioritised variants (weak to empirical), and (iv) genes in the empirical triplets. We retrieved the list of differentially expressed genes (DEGs) for Esophageal Carcinoma (ESCA) versus normal tissues from GEPIA2 and assessed the proportion of DEGs at each level of prioritisation.

Our analysis revealed a significant trend of increasing DEG enrichment with higher TranCi prioritisation. Specifically, the empirical gene set showed a DEG proportion of 16.564%, which was significantly higher than the lower tiers: 13.45% for the weak-to-empirical set (tier iii) and 9.78% for the R2R annotation set (tier ii). The full transcriptome baseline exhibited the lowest proportion at 6.35% (Figure 2d). Two-sample tests for equality of proportions confirmed these differences were statistically significant (Empirical vs Tier ii: p < 2.2e-16; Empirical vs Tier iii: p < 2.2e-16; Empirical vs Tier i: p = 2.38e-7), supporting the biological relevance of high-scoring TranCi target genes in ESCA tumorigenesis. 24 of the DEGs linked to empirical variants were upregulated in tumour samples compared to 4 downregulated DEGs (Figure 2e), highlighting a potential activating role of these regulatory elements.

These findings suggest that genes linked to empirically supported variant–eRNA– mRNA interactions are more frequently dysregulated in ESCA, consistent with the hypothesis that TranCi prioritises regulatory noncoding variants using functional relationships relevant to the disease mechanism.

### Empirical noncoding variant targets enrich for stress, proliferation, and oncogenic signalling pathways

To identify broader molecular and biological themes associated with the empirical variants, we performed gene set enrichment analysis of the 163 associated genes across multiple databases, including Elsevier Pathway Collection, MSigDB Hallmark 2020, Panther 2016, and ClinVar 2019 via the Enrichr platform ^23^. Notably, ethanol-induced hepatotoxicity (Figure 2f) emerged among the top pathways, consistent with the known role of alcohol consumption as a major risk factor in ESCC. Mitotic spindle, a hallmark of cancer-associated proliferation and chromosomal instability, was also among the top hits (Figure 2f). Additional top pathways such as apoptosis signalling, PDGF signalling, Notch signalling, and oxidative stress response further support a cancer-relevant molecular signature (Figure 2f).

Together, these findings suggest that the empirical variant–eRNA–mRNA triplets capture key oncogenic processes, including impaired redox homeostasis and deregulated cell growth signalling, with relevance to ESCC pathophysiology. ClinVar-based enrichment also returned ‘malignant tumour of esophagus’ as the most significant disease term, reinforcing the disease relevance of the evidence (Figure 2g). Although our variant prioritisation focused on noncoding mutations, the enrichment for ESCC-linked coding variants in these same genes suggests a convergence of regulatory noncoding and coding mutations on shared signalling programmes. This coherence points to a broader coordination of transcriptional and post-transcriptional dysregulation in ESCC.

### TranCi-prioritised regulatory axes show functional dependency in ESCC cell lines

To further validate the functional relevance of genes linked to TranCi-prioritised empirical variants, we examined their dependency profiles in ESCC cell lines using CRISPR knockout screen data from the DepMap portal ^24^. This dataset provides Chronos scores, where more negative values indicate stronger gene dependency or essentiality for cell survival and proliferation. We extracted scores across all available ESCC cell lines and classified gene dependency using a threshold-based scheme: scores ≤ -0.5 were considered essential, scores between -0.5 and -0.1 as dependent, and scores > -0.1 as neutral. This classification reflects the biological impact of gene knockout on cell viability, with essential genes being critical for survival.

Our analysis reveals a broad spectrum of dependency among genes linked to TranCi empirical variants (Figure 3a). Notably, 20% (30 out of 146) of these genes demonstrated essentiality in at least one ESCC cell line, highlighting their potential as core drivers. Extending beyond strict essentiality, 80.8% (118 out of 146) were classified as dependent or essential in at least one ESCC cell line, with many dependencies spanning multiple cell lines. These findings support the biological significance of the TranCi-prioritised gene set, corroborating their putative roles in ESCC tumorigenesis and maintenance. They also reinforce the functional relevance of their associated noncoding variants prioritised by TranCi, effectively bridging variant-level predictions with observed cellular phenotypes in cancer models.

**Figure 3.**
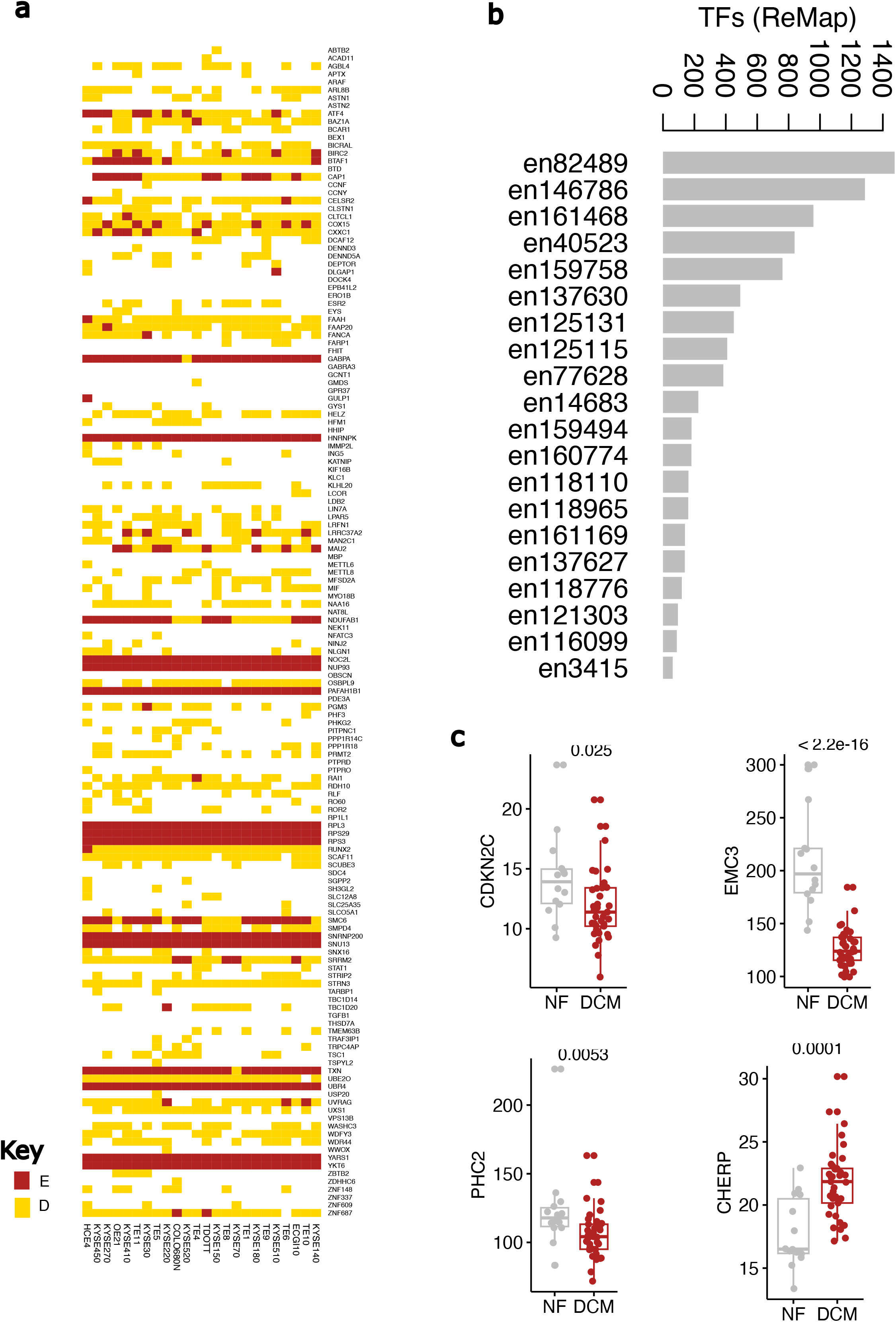
TranCi-prioritised regulatory axes show transcriptional and functional dependency in ESCC and cardiac tissue. (a) Heatmap of CRISPR-Cas9 dependency chronos scores (DepMap 25Q2) for empirical TranCi target genes across ESCC cell lines. Within the plot, red indicate scores ≤ -0.5, essential - E and yellow is scores between -0.5 and -0.1, dependent -D. (b) Number of transcription factors binding to TranCi-prioritised eRNAs based on ReMap 2022 ChIP-Seq database. (c) Boxplots of CDKN2C, EMC3, PHC3, and CHERP expression in non-failing vs. dilated cardiomyopathy (DCM) hearts (GSE116250).

### Widespread TF binding at enhancers limits variant prioritisation by motif analysis

To explore whether transcription factor (TF) binding could aid in functionally annotating these noncoding variants identified by TranCi, we focused on the 20 eRNAs empirically linked to the high scoring variants. We intersected their coordinates with high-confidence TF ChIP-Seq peaks from the ReMap database to quantify TF occupancy. On average, each eRNA region was bound by 475 different TFs, with several overlapping more than 600 TFs (Figure 3b). This extensive co-occupancy severely limits the utility of motif- and TF-centric annotations: with nearly all known TFs binding the same region, it becomes nearly impossible to infer which factor, if any, is functionally relevant to the variant’s effect. These findings underscore the challenge of using TF binding or sequence motifs alone to prioritise noncoding variants.

### Noncoding variant rs11588271 uncovers eRNA–mediated cis–trans regulation of cardiac function

Finally, to demonstrate the utility of TranCi in scenarios where matched RNA expression data is unavailable, we applied the TranCi-explore module. This extension is particularly relevant for interpreting personal genomes or cohorts lacking RNA-Seq, enabling variant annotation through experimental eRNA-RNA interaction map. As an illustrative example, we focused on rs11588271 (chr1:51,004,570), a rare noncoding variant previously associated with QRS duration ^25^, a key cardiac electrophysiological trait.

Upon processing, TranCi-explore identified it as predictive, with a regulatory footprint involving four target genes (CDKN2C, CHERP, EMC3, and PHC2), with en4258, acting as the conduit. Among these, CDKN2C is the only target located proximally and known to reside in the same TAD as en4258 ^19^, supporting direct transcriptional regulation. This aligns with previous eQTL analyses that link rs11588271 to CDKN2C expression, and with our earlier work ^19^, demonstrating that en4258 modulates CDKN2C mRNA stability in a post-transcriptional manner, contributing to phenotypic modulation in vitro. The remaining three genes were implicated through physical eRNA–mRNA interactions. CHERP (Calcium Homeostasis Endoplasmic Reticulum Protein) participates in intracellular calcium release and the calcineurin–NFAT signalling pathway, key mediators of cardiac excitation-contraction coupling and conduction system function ^26^. Its connection to calcium signalling suggests a plausible role in modulating QRS duration via effects on action potential propagation. EMC3, a subunit of the ER membrane protein complex (EMC), facilitates insertion of membrane proteins, including ion channels and GPCRs ^27^. This function may influence cardiomyocyte electrophysiology by regulating membrane protein composition or localisation, thereby affecting ventricular depolarisation. PHC2 (Polyhomeotic Homolog 2), a chromatin-associated transcription repressor, has a less well-defined role in cardiac tissue function but may contribute to heart development or stress-induced remodelling via epigenetic regulation.

To further confirm these assertions, we re-analysed pre-processed gene expression data from normal heart tissue and dilated cardiomyopathy (DCM) ^28^, a condition associated with altered QRS duration. All four genes showed differential expression in DCM: three were significantly downregulated, while CHERP was upregulated (Figure 3c). The three targets linked to rs11588271 exclusively via RNA-RNA interactions showed more pronounced expression changes than CDKN2C. Taken together, these findings expand the mechanistic interpretation of rs11588271, revealing a cis- and trans-acting eRNA-mediated regulatory axis. Therefore, the addition of the TranCi-explore module enables systematic interpretation of noncoding variants in the context of personal genomes, bridging genomic position with regulatory consequence. This extension significantly enhances the translational potential of TranCi, making it more applicable for dissecting molecular mechanisms underlying complex human traits and genetic conditions.

## Discussion

Mapping distal noncoding variants or mutations to their regulatory gene targets remains a major challenge in genomics ^11,12,29^. Although chromatin-based methods and TF motif analysis are commonly used to infer regulatory interactions and impact, these methods often lack specificity. Chromatin contacts can span broad genomic domains, and TF binding to regulatory elements is complex to disentangle.

Moreover, for many variants located outside UTRs, promoters, or regions near known genes, it remains virtually impossible to link TF motifs to their direct downstream targets. This makes it difficult to infer the functional consequences of such variants beyond TF binding alone. TranCi addresses these limitations by integrating eRNA expression and eRNA–mRNA interaction data to identify functionally coherent variant–eRNA–mRNA triplets that represent candidate regulatory axes perturbed by variations. This strategy allows the framework to move beyond genomic proximity, capturing mechanistically supported connections that reflect eRNA-mediated regulatory relationships. By testing for coordinated changes in both eRNA and mRNA expression in variant carriers, coupled with interaction evidence, TranCi enforces a dual-layer filter that enhances specificity and reduces false positives. Unlike existing variant annotation tools, which focus on the enhancer-promoter interactions, TranCi broadens the functional scope of enhancer function to link variants with biological and clinical relevance. This approach builds on emerging evidence that eRNAs are active regulatory molecules with widespread interactions across DNA, RNA, and proteins, enabling precise control of gene expression at the transcriptional, post-transcriptional, and translational levels ^14,15,17,19^.

To address the statistical challenges posed by sparse variant stratification, particularly in cancer genomes where many mutations overlapping eRNA coordinates are rare, TranCi implements an empirical Bayes shrinkage strategy to stabilise variance estimates and enhance power. TranCi borrows information across all transcripts to generate moderated t-statistics ^30^, enabling robust identification of variant-associated eRNA and mRNA changes even in small patient subsets. Beyond expression-based evidence, TranCi leverages RNA–RNA interaction maps to provide mechanistic support for candidate regulatory triplets. These networks, derived from KARR-Seq, RIC-Seq and PARIS, extensively integrated in eRNAkitDB ^18^, allow users to incorporate experimentally observed interactions into the prioritisation framework. While current interaction maps are not tissue-specific to any tissue, their reproducibility across multiple contexts ^17,18^, support their utility in variant-to-function mapping. Future efforts to expand eRNA interaction datasets in a tissue- and disease-specific manner will further improve this aspect of the framework.

To assess the biological relevance of TranCi-prioritised triplets in ESCC, we integrated gene expression, pathway enrichment, disease annotation, and gene dependency data. Predicted target genes were consistently altered in ESCC tumour samples relative to normal tissue and enriched for canonical oncogenic pathways, including apoptosis signalling, mitotic spindle regulation, and alcohol-related risk factors. Many targets also overlapped with genes implicated in esophageal malignancy and genomic instability syndromes, and most of these targets exhibited tumour essentiality and dependency in CRISPR knockout screens. These findings support the hypothesis that noncoding mutations modulating eRNA levels may drive tumour phenotypes by rewiring gene expression programs through RNA-mediated regulation. Notably, TranCi avoids reliance on motif-based annotations or TF binding models, which suffer from significant redundancy. On average, enhancer regions linked to the associated eRNAs in ESCC were bound by over 475 TFs, with some exceeding 600. This extensive co-occupancy renders motif-centric analyses uninformative for prioritising specific regulatory drivers. In contrast, TranCi leverages transcriptomic and interaction data to infer functional effects, sidestepping the ambiguity of TF-based inference in enhancer-dense regulatory regions.

While TranCi demonstrates significant advantages, several limitations merit discussion. First, the framework is constrained by the completeness and context-specificity of current RNA–RNA interaction datasets, which are not fully representative of all cell types and conditions. However, the mechanistic reproducibility of these interactions, along with their validation in multiple models, suggests that this limitation can be mitigated, particularly through the large volume of data curated in eRNAkitDB. Additionally, the use of empirical evidence from the input samples further alleviates this issue by refining interactions directly from expression data. Nonetheless, this expression-based approach restricts the analysis to enhancers with detectable eRNA transcription, meaning that non-transcribed or constitutively degraded but potentially functional regulatory elements are not captured. There is scope to expand the approach by incorporating nascent RNA measurements, such as PRO-Seq or GRO-Seq, which may identify additional transient or low-abundance eRNAs. However, these data types are not routinely matched with WGS datasets and often suffer from sparsity and noise, limiting their practical application. Despite this, the current version of TranCi leverages a comprehensive set of over 45,000 transcribed and detectable enhancer annotations, providing a strong foundation for mapping regulatory interactions. Furthermore, the current version of TranCi does not explicitly incorporate RNA secondary structure or RNA-binding protein recruitment, although these aspects are under active development in ongoing work.

To enable broader application, we developed TranCi-explore, a single-sample mode that allows users to interrogate variant function in individual genomes. This personalised analysis mode is particularly relevant for clinical settings, where rare or patient-specific noncoding variants or mutations must be interpreted in isolation.

While not the primary focus of this manuscript, TranCi-explore represents a powerful extension of the framework, and the use case presented here demonstrates its clinical utility using a known functional variant.

In summary, TranCi provides a biologically informed and statistically robust framework for interpreting noncoding variants across a wide range of contexts. By integrating eRNA expression, variant data, and RNA–RNA interaction maps, it enables mechanistically grounded variant prioritisation that goes beyond traditional annotation methods. The approach is particularly well-suited to cancer and rare disease applications, where regulatory mutations are common yet challenging to functionally characterise. With a modular architecture and expanding data resources, TranCi offers a scalable solution for linking noncoding variation to gene regulation and disease biology.

## Method

### TranCi Framework

We developed the TranCi framework to prioritise functional non-coding variants by leveraging enhancer RNA (eRNA)–mediated post-transcriptional interactions as a mechanistic link between non-coding variants and gene function (Figure 1). Our approach draws on previous efforts ^11,12,29^ to annotate non-coding variants in the human genome but advances chromatin-centric models by incorporating post-transcriptional eRNA regulatory interactions. Specifically, while existing tools define the regulatory potential of noncoding variants based on chromatin accessibility, histone modifications, or transcription factor motifs, functional eRNA–mRNA and eRNA–protein interactions ^14,15,17,19^ provide an orthogonal layer of evidence that reveals downstream regulatory consequences, including effects on mRNA stability, localisation, and translation. Together, these layers help uncover regulatory mechanisms not captured by transcription-centric annotations alone.

We posited that post-transcriptional interactions between eRNAs and their target mRNAs offer a more direct and functionally informative route for linking non-coding variants to gene regulation. Based on this assumption, we modelled variant-induced regulatory cascades as originating from changes in eRNA expression, driven by variants overlapping eRNA loci, and propagating through eRNA–mRNA interactions to influence target gene expression and function.

TranCi begins by identifying genetic variants that overlap the genomic coordinates of expressed eRNAs. The framework takes four primary inputs: (i) a path to VCF files containing variant calls across the cohort, (ii) an expression matrix of eRNAs, (iii) an expression matrix of mRNAs, and (iv) a reference annotation database from eRNAkit. For each variant, TranCi stratifies the cohort based on genotype and tests for differential eRNA expression between carriers and non-carriers using empirical Bayes moderated t-statistics, implemented using an adapted *limma*-based approach ^30^. To account for small carrier group sizes, variance estimates are stabilised by borrowing information across all eRNAs through global empirical Bayes priors ^30–32^. After identifying eRNAs significantly associated with genetic variants, the framework then retrieves their putative target mRNAs based on RNA–RNA interaction pairs from the supplied eRNAkit reference database. TranCi tests whether these target mRNAs also exhibit differential expression between variant carriers and non-carriers. The joint evidence of coordinated expression changes in both eRNAs and their interacting mRNAs forms the basis of the empirical support for functional variant impact.

In parallel, TranCi annotates all eRNA–mRNA interaction pairs with predictive features related to post-transcriptional regulation, including impact on target mRNA stability and translation efficiency. These annotations, sourced from ribosome profiling, decay assays, and subcellular localisation data integrated in eRNAkit, are applied regardless of whether the mRNA target exhibits expression changes. This allows TranCi to identify potential regulatory consequences even in the absence of measurable mRNA shifts, capturing additional layers of functional impact.

### Empirical Bayes Moderation of Differential Expression Statistics

Given that variants overlapping eRNA loci are often rare within cohorts, sometimes represented by as few as two samples (TranCi minimum threshold = 2), TranCi employs an empirical Bayes framework to stabilise variance estimates in differential expression testing of both eRNAs and mRNAs between carriers and non-carriers.

This approach borrows information across the entire expression dataset to improve variance estimation, enhance statistical power, and reduce false discovery rates in genotype-stratified analyses with sparse sample groups. The method proceeds in two key steps. First, a global variance prior is estimated by fitting a simple linear model with only an intercept term to all expression data, capturing background variability common across genes or eRNAs. Applying the empirical Bayes method ^30^ to this global fit yields hyperparameters including a global prior variance estimate and prior degrees of freedom. Second, for each variant-associated subset of expression data, TranCi fits a linear model contrasting variant carriers versus non-carriers. Residual variances from these models are combined with the global prior variance in a weighted manner to produce posterior variance estimates, defined as:

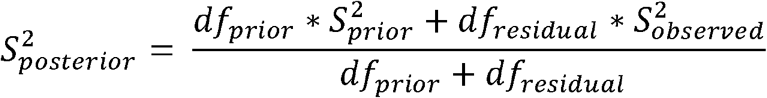

Where, S is variance estimate and df is degree of freedom.

This shrinkage of noisy variances towards the global prior stabilises variance estimates, which is critical when carrier numbers are low. Using these posterior variances, moderated t-statistics are computed for the genotype effect, with degrees of freedom adjusted accordingly. Two-sided p-values are calculated and corrected for multiple testing using the Benjamini-Hochberg procedure to control false discovery rates.

### Integrative Scoring of eRNA–mRNA Pairs to Quantify Variant Impact

To quantify the strength of evidence supporting each variant-eRNA–mRNA interaction, an integrative score was assigned to every pair based on a weighted sum of mechanistic and expression-based indicators. All interactions began with a baseline score of 0.5, reflecting the premise that any variant-eRNA–mRNA interaction suggests a potential regulatory effect. Additional points were added depending on whether key features passed a significance threshold. Specifically, the score increased by 5 if the eRNA showed statistically significant of (p < 0.05) differential expression between variant carriers and non-carriers, by 2.5 if the target mRNA was also significantly differentially expressed in the same group (FDR < 0.1), by 1 if there was evidence that the eRNA modulate the stability of the target mRNA and by 1 if eRNA modulate translational efficiency of the target mRNA (FDR < 0.1). These weighted increments prioritised direct transcriptional and downstream regulatory consequences while still allowing for intermediate or partial evidence to contribute to the overall interaction score. Note: The current implementation of TranCi is limited to single nucleotide variants or mutations. As such, eRNA effects are called using raw p-values, since only one eRNA annotation can overlap each variant. Feature modules could expand this to include other variant types, such as insertions, deletions, or structural variants, enabling broader assessment of eRNA regulatory effects.

To consolidate evidence across multiple eRNA–mRNA interactions linked to the same variant, a cascading approach was used to derive a single variant-level score. For each variant, the maximum score among all associated eRNA–mRNA pairs was selected as its representative score. This strategy ensured that a strong mechanistic signal from even one interaction was sufficient to elevate the variant’s prioritisation, rather than requiring uniform support across all linked transcripts. Variants were then categorised into three tiers based on their maximum interaction score: those with scores of 6.5 or higher were classified as “empirical,” indicating robust support from both eRNA and target expression data; those with scores between 1.5 and 6.5 were considered “predictive,” reflecting partial but meaningful support; and those with scores below 1.5 were labelled “weak,” indicating limited or inconclusive evidence.

This tiered scoring framework allowed for systematic triaging of variant candidates based on the strength of inferred enhancer-mediated regulation.

### TranCi-explore: Variant Prioritisation in the Absence of RNA-Seq

While the core TranCi framework leverages matched total RNA-Seq and genotype data to identify noncoding variants with empirical evidence of post-transcriptional regulation, such datasets are not always available, particularly in clinical or personal genomics settings. In these contexts, users often have access to variant data (e.g. VCF files from whole genome, coordinates or bed files) but lack matched transcriptomic measurements. To address this limitation, we developed TranCi-explore, a complementary mode designed to prioritise personal or rare noncoding variants using only genotype information.

TranCi-explore operates by systematically mapping variants to annotated eRNA loci and evaluating their potential regulatory impact through reference knowledge derived from large-scale functional datasets. Specifically, it links each variant to nearby eRNAs and retrieves all known eRNA–mRNA interaction pairs from the eRNAkit database. These interactions are then annotated using predictive features, including known effects on mRNA stability, translation efficiency, and subcellular localisation (same as the core module), providing a mechanistic basis for variant prioritisation even in the absence of RNA expression data. Unlike the cohort-based mode, TranCi-explore does not rely on differential expression testing but instead uses precomputed annotations and reference models to assign scores based on the presence and nature of regulatory interactions.

## Datasets

### eRNA Annotation and RNA-RNA interaction curation

The eRNA annotation across multiple cell types and the compilation of high-confidence RNA-RNA interactions (from KARR-Seq, RIC-Seq, and PARIS datasets) and eRNA impact on mRNA stability and translation efficiency used in this study were previously described in ^18,19^.

### ESCC samples

We obtained TCGA data via the NCI Genomic Data Commons (GDC) Data Portal, which provides uniformly processed and harmonized datasets aligned to GRCh38 (hg38) using standardised bioinformatics pipelines. Somatic mutation calls (VCF), RNA⍰ Seq aligned BAMs, were accessed through the GDC portal with dbGaP authorisation (phs000178/GRU). Preprocessing of RNA-Seq BAM files to quantify eRNA and mRNAs were performed using our established pipeline ^18^.

### Heart samples

We retrieved normalized RNA-Seq expression data (RPKM) for non-failing and dilated cardiomyopathy heart samples from GEO accession GSE116250 ^28^. The processed RPKM expression matrix, as provided by the original authors, was directly downloaded from the GEO Omnibus repository and used for downstream analysis without further transformation.

### DepMap

Gene dependency data for ESCC cell lines were obtained from the DepMap portal (https://depmap.org/portal), release 25Q2. CERES-corrected CRISPR dependency scores were used to assess the functional relevance of empirical eRNA-associated genes across ESCC models. Cell lines with dependency scores below −0.5 were considered to exhibit essentiality, those between -0.5 and -0.2 were considered dependent.

### Functional annotation and enrichment analysis using online platforms

RegulomeDB v2.2 was used to assess the regulatory potential of each variant. The search was performed via the online interface (https://regulomedb.org/) using default settings. Variants were categorised based on assigned functional classes and associated probabilities.

DeepSEA predictions were obtained via the online platform (https://deepsea.princeton.edu/, new version) to estimate regulatory impact scores for each variant. We extracted per feature scores and summed the probabilities to create composite regulatory potential score.

For gene-level functional interpretation, Enrichr (https://maayanlab.cloud/Enrichr/) was used to perform gene set enrichment analysis on empirical eRNA-associated genes. Enrichments for Elsevier Pathway Collection, MSigDB Hallmark 2020, Panther 2016, and ClinVar 2019 were retained for visualisation and interpretation.

## Statistical analysis

Unless otherwise stated, graphical data represent the mean ± standard error of the mean (SEM) from at least three independent experiments. Differences between groups were assessed using Wilcoxon rank-sum test. Statistical significance was defined as p□ <□ 0.05 for single comparisons.

## Data availability

The TranCi module is provided as part of the eRNAkit R package, available at https://github.com/AneneLab/eRNAkit. This toolkit, described in a separate work ^18^, includes the eRNAkitDB used to power TranCi. Prioritised variant-eRNA-gene triplets and noncoding variant classes are also included within the supplemental table.

## Author contributions

Project was conceived of by C.A.A. Data curation and processing were done by C.A.A, N.B, and R.K. Analysis and interpretation was done by C.A.A, J.H, J.B, N.B, R.K, C.A.A designed the TranCi framework and wrote the manuscript. All authors edited the final manuscript.

